# Dairy vs. beef production – expert views on welfare

**DOI:** 10.1101/2021.12.06.471462

**Authors:** Roi Mandel, Marc B.M. Bracke, Christine J. Nicol, John A. Webster, Lorenz Gygax

## Abstract

Consumers’ views and concerns about the welfare of farm animals may play an important role in their decision to consume dairy, meat and/or plants as their primary protein source. As animals are killed prematurely in both dairy and beef industries, it is important to quantify and compare welfare compromise in these two sectors before the point of death. Seventy world-leading bovine welfare experts based in 23 countries, were asked to evaluate the likelihood of a bovine to experience 12 states of potential welfare concern, inspired by the Welfare Quality® protocol. The evaluation focused on the most common beef and dairy production systems in the experts’ country, and was carried out separately for dairy/beef calves raised for red-meat, dairy/beef calves raised for veal, dairy/beef calves raised as replacement, and for dairy/beef cows. The results show experts rated the overall likelihood of a negative welfare state (i.e. welfare risk) to be higher in animals from dairy herds than from beef herds, for all animal categories, regardless of whether they were used to produce milk, red-meat or veal. These findings suggest that consuming food products derived from common dairy production systems (dairy or meat), may be more harmful to the welfare of animals than consuming products derived from common beef production systems (i.e. from animals solely raised for their meat). Raising awareness about the linkage between dairy and meat production, and the toll of milk production on the welfare state of animals in the dairy industry, may encourage a more sustainable and responsible food consumption.

## Introduction

Abstaining from consuming meat, but not dairy products, is regarded by many to be a sign of compassion towards animals [1,2]. Vegans go a step further and reject the consumption of any animal-based product, but few, if any, advocate abstention from dairy while continuing to consume meat. The extent to which an independent analysis of animal welfare aligns with such consumer choices has not previously been examined. Importantly, there are substantial reasons to predict a discrepancy, particularly when considering the role of the dairy industry in producing meat (Figure 1A), and the higher degree of intervention in the lives of animals that are used “for more than their meat” (i.e. intervention associated with daily milking of the dams, and the consequent management of their calves). Here, we obtained an independent assessment of the likelihood that dairy cattle and beef cattle would experience negative welfare, by surveying a panel of leading cattle welfare experts, each focusing on the most common beef and dairy production systems in their country of expertise. The comparison was carried out between dairy and beef calves when raised directly for their meat (i.e. dairy and beef calves raised for red-meat and for veal, consisting mostly of male calves), and between dairy and beef cows raised for producing calves/milk.

**Figure 1.**
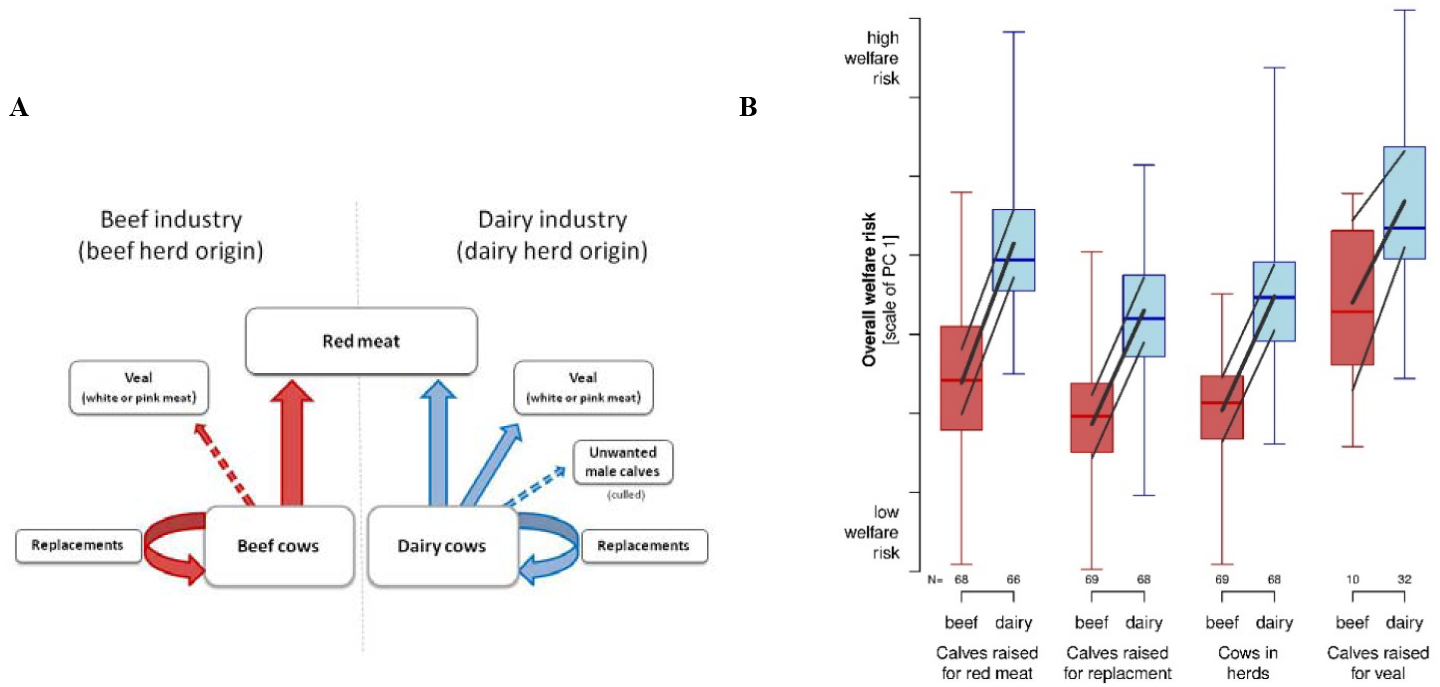
**(A)** Diagram showing the origin (dairy/beef herd) and production goals, i.e. the general flow of calves from dairy herds and beef herds to red-meat, veal and cow replacements (inspired by EFSA Panel on Animal Health and Welfare [19]. Dashed arrows reflect production routes that are not common to all farms. **(B)** First principal component reflecting the overall welfare risk (likelihood of 12 welfare concerns, see text) as assessed by the experts as a function of the different origins and animal production goals. N: Number of experts that reported an assessment on all 12 areas of welfare concern. Boxplots: show minimum, lower quartile, median, upper quartile and maximum values. Black lines: model estimates with 95% upper and lower confidence intervals.

Expert assessments can help to characterise uncertainty and fill data gaps where traditional scientific research is not possible or data are not yet accessible or available [3]. Such assessments have been widely used to review impacts of housing, management or other anthropogenic challenges to domestic (e.g. [4] cattle; [5] broilers and [6] canine and felines) and wild animals (e.g. [7]), using frameworks such as the Five Domains (e.g. [7,8]), and more recently, in relation to the United Nations Sustainable Development Goals [9,10]. The European Food and Safety Authority (EFSA) commonly uses expert opinion to inform debates around welfare topics, such as the use of perches for laying hens [11], and for assessing welfare risks, such as those that relate to the farming of sheep for wool, meat and milk production [12].

In the next section, we compare experts’ assessment of the overall likelihood that bovines will experience a negative welfare state when raised in the most common housing systems in the dairy and beef sectors, with a focus on Europe and North America. We predicted that the higher degree of intervention in the lives of dairy cattle, which stems from the fact that they are used for “more than their meat” (i.e. harvesting of milk, affecting their management and the management of their calves), would result in dairy cattle being rated by experts as being at a higher welfare risk than beef cattle.

## Methods

An approval to conduct this study was granted by the Research Ethics Committee for SCIENCE and HEALTH of the university of Copenhagen, ref: SUSTA, case: 504-0272/21-5000. Approval and registration of processing of personal data in the project was provided by the legal department of the university of Copenhagen, ref PWI, case: 514-0273/21 – 5000.

### Data Collection

Overall, 130 cattle welfare experts were invited to participate in our survey. based on their 1. Number of publications on the topic of bovine welfare (peer reviewed research manuscripts and review articles) that appear in the Web of Science data base, when using the following key words: “Welfare” + “bovine”/”cattle”/”dairy cow”/”beef cow”) (selected experts were those with the highest number of publications) and 2. H-index (minimum of 10). In addition, we invited further researchers that were recommended to us by the selected experts. Overall, 83 experts agreed to participate in the survey, out of which 13 were omitted from the final analysis for the following reasons: 10 experts did not complete the survey, 2 experts felt that they could not provide accurate assessments, and one expert felt that his degree of expertise was not sufficient. Data were collected through an online survey built in Qualtrics (Qualtrics XM Platform™, Provo, Utah, USA), between October and December 2020.

The survey included four parts: 1. Consent including a short description of the aim of the experiment “to compare the welfare of cattle across food production systems”. 2. General instructions. 3. Characterization of the common production systems, likelihood ratings and confidence ratings. 4. Characterization of experts (as summarized in S1 Table).

#### Characterization of the common production systems

Before providing likelihood ratings, the experts were asked to describe the most common beef and dairy production systems in their country of expertise. The characterization applied to each animal category separately, using 3-5 fixed criteria (S1 Fig.; S3 Table), to which the experts could answer: “yes”, “partial/part of the time”, “no”, or “I don’t know”.

#### Likelihood rating

The experts were asked to rate the likelihood of 12 statements on a scale of 1 (very low) to 7 (very high), and to notice that the statements were built in such a way that the higher the likelihood ratings are, the higher is the welfare-risk for the animal (for illustration see S1 Fig., for the exact questions and ratings see Table 1). The 12 short welfare statements that were inspired by the Welfare Quality protocol [13], a well-established protocol for assessing bovine welfare. The statements addressed the following core areas of potential welfare concern: 1. inadequate diet, 2. inadequate water supply, 3. thermal discomfort, 4. resting discomfort, 5. injuries, 6. disease, 7. pain resulting from management/handling/surgical procedures, 8. inability to move freely, 9. inability to perform social behavior, 10. inability to perform other normal behaviors, 11. experiencing negative affective states, and 12. lack of experiencing positive affective states (for the order of presentation, see Table 1). Since management practices were expected to vary between countries for both beef and dairy sectors (for red-meat, veal, replacement, and cows), animal categories were defined as follows: For red meat calves and veal, the evaluation period was from birth to slaughter (or up to 18 months of age). For replacement calves it was from birth to first calving. For cows, it was from first calving to slaughter. In all cases, the evaluation did not include transportation to slaughter or the process of slaughter itself. To avoid order effects (where rating one animal origin first (e.g. beef) would affect the rating of the other (e.g. dairy), we used 4 complementary versions of the survey. In all four versions, the presentation of animal category was fixed: Red-meat, replacement calves, cows and veal. However, the order of presenting the animal origin (beef/dairy) was counter balanced (left/right*text colour blue/orange). In each of the four versions, the text colour and order of presentation in Fig. 1A was matched accordingly.

**Table 1.** likelihood – likelihood and confidence ratings for each of the 12 statements. Raw data. Mean (STD) and median [min – max].

| Animal category<br>Animal origin (herd) | Red-meat |  | Replacement Calves |  | Cows |  | Veal |  |
| --- | --- | --- | --- | --- | --- | --- | --- | --- |
|  | Beef | Dairy | Beef | Dairy | Beef | Dairy | Beef | Dairy |
| **Q1:** Receive inappropriate/inadequate diet (e.g. quantity of feed, number of feeding bouts, quality of feed, digestibility etc.). | 2.4 (1.4)<br>2 [1-7] | 4.2 (1.9)<br>4 [1-7] | 2.2 (1.4)<br>2 [1-7] | 3.4 (1.7)<br>3 [1-7] | 2.8 (1.3)<br>3 [1-7] | 2.7 (1.6)<br>2 [1-7] | 3.5 (2.0)<br>4 [1-7] | 4.7 (1.9)<br>5 [1-7] |
| **Q2:** Have inadequate water supply (e.g. quantity, quality, accessibility, high competition over access to water). | 2.2 (1.3)<br>2 [1-7] | 2.6 (1.5)<br>2 [1-7] | 2.0 (1.2)<br>2 [1-7] | 2.4 (1.5)<br>2 [1-7] | 2.4 (1.3)<br>2 [1-7] | 2.3 (1.4)<br>2 [1-7] | 3.3 (2.3)<br>3 [1-7] | 3.4 (1.8)<br>4 [1-7] |
| **Q3:** Experience discomfort when resting (e.g. when lying down, standing up etc.). | 2.4 (1.6)<br>2 [1-7] | 3.8 (1.7)<br>4 [1-7] | 2.1 (1.3)<br>2 [1-7] | 3.1 (1.5)<br>3 [1-7] | 2.2 (1.3)<br>2 [1-6] | 3.7 (1.5)<br>4 [1-7] | 3.6 (2.3)<br>4 [1-7] | 4.5 (1.6)<br>5 [1-7] |
| **Q4:** Experience thermal discomfort (i.e. being too warm (e.g. heat load) or too cold). | 3.3 (1.4)<br>3 [1-7] | 3.3 (1.6)<br>3 [1-7] | 3.0 (1.4)<br>3 [1-6] | 3.1 (1.4)<br>3 [1-6] | 3.2 (1.4)<br>3 [1-6] | 3.3 (1.5)<br>3 [1-7] | 3.7 (1.6)<br>4 [2-6] | 3.9 (1.5)<br>4 [1-6] |
| **Q5:** Experience restricted movement (e.g. limited ability to move around freely, e.g. being tied, kept in high stocking density). | 2.4 (1.6)<br>2 [1-7] | 4.3 (1.7)<br>4 [1-7] | 2.0 (1.3)<br>2 [1-7] | 3.6 (1.5)<br>3 [1-7] | 2.0 (1.2)<br>2 [1-7] | 3.6 (1.7)<br>4 [1-7] | 3.5 (2.1)<br>4 [1-7] | 5.0 (1.5)<br>5 [2-7] |
| **Q6:** Suffer from injuries (e.g. lameness, skin lesions etc.). | 2.6 (1.4)<br>2 [1-7] | 3.4 (1.5)<br>3 [1-7] | 2.2 (1.1)<br>2 [1-7] | 2.9 (1.3)<br>3 [1-7] | 2.7 (1.3)<br>3 [1-7] | 4.7 (1.4)<br>5 [2-7] | 3.0 (1.8)<br>3 [1-6] | 3.9 (1.5)<br>4 [1-7] |
| **Q7:** Suffer from disease | 3.1 (1.4)<br>3 [1-6] | 4.2 (1.6)<br>4 [1-7] | 2.6 (1.1)<br>2 [1-7] | 3.6 (1.4)<br>4 [1-7] | 2.7 (1.1)<br>3 [1-6] | 4.2 (1.6)<br>4 [1-7] | 3.8 (1.8)<br>4 [2-6] | 5.0 (1.4)<br>5 [1-7] |
| **Q8:** Suffer from pain induced by management, handling, or surgical procedures | 4.1 (2.0)<br>4 [1-7] | 4.4 (1.8)<br>4 [1-7] | 3.2 (1.8)<br>3 [1-7] | 4.1 (1.9)<br>4 [1-7] | 2.7 (1.4)<br>2 [1-7] | 3.4 (1.6)<br>3 [1-7] | 3.5 (1.8)<br>3 [2-7] | 3.8 (1.7)<br>3 [1-7] |
| **Q9:** Have limited opportunities to express normal, non-harmful, social behaviors (e.g. limited allo-grooming, limited social play etc.). | 2.2 (1.5)<br>2 [1-7] | 4.0 (1.9)<br>4 [1-7] | 2.0 (1.2)<br>2 [1-7] | 3.4 (1.7)<br>3 [1-7] | 1.9 (1.1)<br>2 [1-6] | 3.2 (1.4)<br>3 [1-6] | 3.8 (2.2)<br>4 [1-7] | 5.0 (1.5)<br>5 [2-7] |
| **Q10:** Have limited opportunities to express other normal behaviors (i.e. limited/restricted from opportunity to forage/graze). | 2.7 (1.7)<br>2 [1-7] | 4.9 (1.7)<br>6 [1-7] | 1.9 (1.2)<br>2 [1-6] | 3.9 (1.7)<br>4 [1-7] | 1.8 (1.2)<br>2 [1-7] | 3.9 (1.5)<br>4 [1-7] | 4.1 (2.4)<br>4 [1-7] | 6.0 (1.3)<br>6 [2-7] |
| **Q11:** Experience negative affective states (e.g. fear, distress, frustration or apathy; due to negative human-animal relationships, husbandry / management or the environment in which the animal is kept). | 3.0 (1.5)<br>3 [1-7] | 4.3 (1.7)<br>4 [1-7] | 2.6 (1.2)<br>2 [1-7] | 3.4 (1.5)<br>3 [1-7] | 2.5 (1.3)<br>2 [1-7] | 3.6 (1.6)<br>4 [1-7] | 4.2 (2.3)<br>4 [1-7] | 4.9 (1.7)<br>5 [1-7] |
| **Q12:** Have limited opportunities to experience positive emotions (e.g. limited opportunities to engage in rewarding activities). | 2.7 (1.6)<br>2 [1-7] | 4.5 (1.6)<br>5 [1-7] | 2.2 (1.3)<br>2 [1-6] | 3.6 (1.5)<br>4 [1-7] | 2.2 (1.2)<br>2 [1-6] | 3.6 (1.4)<br>4 [1-6] | 3.6 (2.3)<br>3 [1-7] | 5.1 (1.7)<br>5 [1-7] |
| **Confidence in ratings** | 3.9 (0.9)<br>4 [1-5] | 3.8 (0.8)<br>4 [2-5] | 3.8 (1.1)<br>4 [1-5] | 4.1 (0.8)<br>4 [2-5] | 4.0 (1.0)<br>4 [1-5] | 4.3 (0.7)<br>4 [2-5] | 3.9 (1.0)<br>4 [2-5] | 3.8 (1.0)<br>4 [1-5] |

#### Confidence rating

The experts were asked to state their level of confidence in their likelihood ratings on a 5 point scale (1 star = low confidence to 5 stars= high confidence). Confidence ratings were reported separately for each of the animal origins in the four animal categories (S2 Fig., and Table 1).

### Data processing

We omitted all answers from respondents that had provided an incomplete set of answers for a given combination of animal origin and category. This left the answers of 70 experts (out of 83). In total 1, 3, 34, 25, and 7 experts gave complete answers for 3, 4, 6, 7, and 8 of the 8 possible combinations of origin and animal category, respectively.

#### Calculating normally-distributed values based on the likelihood ratings

The likelihood ratings were reported on a seven point Likert-scale. A Likert-scale has a fixed lower and upper limit and is therefore comparable to a proportion. Accordingly, it can be expected that a logit-transformation will lead to normally distributed data points. Two experts indicated an interval for the Likert-scale ranging one scale point instead of a single number. In these cases, we used the mid-point of the interval (i.e. the average of the two indicated scale points), which resulted in non-integer scale points at half the original points. Consequently, the resulting Likert-scale had 13 scale points: the integers from 1 to 7 and all the midpoints between the integers (1.5, 2.5, …). In a next step, we re-scaled the 13-point Likert scale to a score on a scale from 0 to 1. We did this in a way that kept the steps between the points of the Likert-scale equal and shifted the minimum and maximum scale points half the distance between the points away from 0 and 1: proportion = 4/(2*13) Likert-scale – 3/(2*13) (S3 Fig.). We then logit transformed the numbers of this 0-1 scale (“normalised values”).

The likelihood ratings were dependent on expert who rated the likelihood of the 12 welfare statements once for each of the two origins and for the four animal categories. To account for this dependency, we calculated the residuals of a mixed model with expert ID as the sole random effect and the intercept as the fixed effect using the 12 normalised values as an outcome variable each. In doing so, we adjusted the average likelihood rating of each expert to the overall average of all experts. Quantile-quantile plots of these 12 sets of residuals showed that they were very close to normally distributed. They thus provided the data for running a principal component analysis (PCA).

### Statistical analysis

For statistical evaluations R V 4.0.3 was used [14]. A PCA based on the correlation matrix (function princomp; base R) was run for each set of variables that reflected information on the most common housing systems of the different animal categories in the countries for which experts made an assessment. These PCAs were based on raw scorings of the answers (no= 1, partly= 2, yes= 3) and responses of experts with missing answers were omitted. Veal from beef origin was also omitted because information for this category was given by less than 15% of the experts (n = 10 of 70).

The residuals of normalised values for the 12 welfare statements (see above) were subjected also to a PCA. The first principle component of this PCA, which explained 50.3% of the total variance, was then used as the outcome variable in a linear mixed-effects model (function blmer, package blme [15], which is based on lme 4 [16]). The fixed effects in the model were the origin of the animals (two-level factor: dairy versus beef), the animal category (four-level factor: calves raised for red-meat, calves raised for replacement, cows, and veal calves) and their interaction. To be able to interpret average main effects even in the presence of interactions, sum-contrasts were used for the fixed effects. The random effect was animal category nested in expert and the confidence ratings were used as weights. Residuals were checked graphically, using a QQ-plot of the raw residuals and the standard plots using simulated residuals in package DHARMa [17]. The model with the confidence weights showed a slight s-shape in the QQ-plot of DHARMa but the estimated effects were almost unchanged in comparison with the model without weights. We therefore report the model with weights. We calculated p-values by comparing the maximum model with one model each that omitted one of the fixed effects using a parametric bootstrap (package pbkrtest [18]). A few warnings about non-convergence occurred in the process of evaluation but these did not seem to influence the model estimates in any relevant way. Finally, we also used a parametric bootstrap to estimate confidence intervals of the model estimates, which fitted the raw data well (Fig. 1).

## Results

Overall 70 cattle welfare experts participated in the survey. The experts, who had a median experience of at least 15 years, were recruited from Europe (36), North America (17), South America (8), Australia (5), and other regions of the world (4). Additional characteristics of the experts can be found in S1 Table.

Experts’ likelihood and confidence ratings for each of the 12 statements are shown in Table 1. The overall likelihood to experience a negative welfare state (i.e. welfare risk) was assessed by the experts as higher in animals from dairy in comparison to beef, for all animal categories (origin: p = 0.001, interaction: p = 0.28; Figure 1B). The overall welfare risk increased from calves raised for replacement and cows, to calves raised for red-meat and to veal calves (p = 0.001; Figure 1B). These results indicate that, regardless of the production goal (calves raised for red-meat, calves raised for veal, calves raised as replacement, and cows), animals born in dairy herds were considered to experience worse welfare than animals born in beef herds.

## Discussion

“Harvesting” milk from bovines has a long tradition (“secondary products revolution” [20]), and it seems to have a promising future, based on recent projections by the OECD as well as the FAO [21]. For this reason, it is important to critically reflect on the welfare impact of this practice. Our expert survey showed that bovines raised in main current food production systems are rated as more likely to experience negative welfare conditions if raised in dairy systems (regardless of whether they are used for their milk or meat) than in beef systems (i.e. when raised solely for their meat). This result may be surprising in light of societal perceptions regarding ethical food choices (vegetarianism [1,2]) and deserves further consideration to identify candidate underpinning reasons.

Our assessment protocol allowed us to cover a wide range of production methods (for the result of the PCAs for the information on the housing systems, see S5 Fig.) and to assess experts ratings of the overall welfare risk of animals raised in these conditions during the majority of their life-time (for the ‘raw’ ratings of experts, for each of the 12 areas of welfare concern, see Table 1). Nevertheless, it did not allow us to disentangle which of the welfare risks mentioned above formed the basis for the experts’ ratings. In addition, we emphasize that the results do not necessarily mean that animals born in dairy herds are, at any given point of time and in every type of system, worse off than animals born in beef herds. For example, after spending several months on pasture with their dam, the welfare of beef suckler calves is expected to be substantially reduced once they are moved to feedlots, potentially below the levels experienced by many dairy calves raised for red-meat (see Figure 1 [35]; for a review of the factors affecting cattle welfare in feedlots see [36,37]; for cattle preferences to being on pasture over the feedlot area see [38]). It would be interesting to explore whether experts would consider that improvements to the welfare of bovines in the dairy industry e.g. by keeping dairy calves with their dams [39], free access to pasture [40] and the use of pain relief, e.g. in case of lameness [41] and during practices such as dehorning [42] would be sufficient to balance the overall welfare risk for animals raised in the two production systems. The answer cannot be predicted in advance because, for example, if such measures were similarly applied to bovines in the beef industry, then they may retain their perceived higher welfare status.

One possible reason why experts rated dairy systems as more harmful to welfare may be because harvesting milk from bovines involves a higher (negative) intervention in their lives compared to raising them solely for their meat Raising bovines for their meat involves feeding and slaughtering them. Dairy cows, are also bred for their milk, which is then collected 1-3 times per day, often for 305 days or more per lactation [22,23], and this has implications for how these animals are raised and managed. Long term genetic selection for high milk yield in dairy cows has been recognized as a major factor causing poor welfare, in particular health problems, such as lameness, mastitis, reproductive disorders and metabolic disorders [24]. Adoption of high yielding breeds such as the Holstein-Friesian without consideration for the animals’ natural ability to cope with diseases [24] and thermal challenges typical of extreme climates may lead to additional welfare compromises [25,26]. Common housing and husbandry procedures characterizing the management of dairy cattle increase the welfare risks further. For example, dairy calves, in contrast to beef calves, are commonly separated from their dam a few hours after birth to allow the collection of milk from their mothers [24,27]. Upon separation, they are often kept in social isolation for several weeks (i.e. limited or no physical contact with their dam or conspecifics), a management practice that originated from the desire to reduce horizontal transmission of disease between calves, yet has been repeatedly shown to inflict behavioural and developmental harm [28]. The social and nutritional restrictions following the separation from the dam have been implicated as key animal welfare issues in commercially raised dairy calves [29], with the former being associated with cognitive and social impairments effecting the welfare of the animals also at a later age. Male dairy calves are often transported to a different farm at a young age, increasing their welfare risk further [30]. Later in life, dairy calves raised to produce milk (“replacement dairy calves”) are commonly housed indoors for part of the year (i.e. winter) or continuously (i.e. zero grazing system), in free or tie-stall systems [31]. Keeping dairy cattle indoors is associated with numerous behavioural restrictions [32,33] and health risks, such as higher incidence of lameness [34], and increased risk for claw or foot problems, teat trampling, mastitis, metritis, dystocia, ketosis, retained placenta, and some bacterial infections compared to systems that allow cattle access outdoors [24]. Since dairy cattle are commonly raised indoors for at least part of the year, and since their management involves a high degree of intervention as described above (in the first weeks of life following the separation from their dam, and throughout the lactating periods, when milked by partially/fully automated milking systems), their management is expected to involve a higher risk to their welfare compared to beef cattle. A future study could specifically explore which of these factors most influence expert rankings of welfare.

Our study focused on the welfare risks of dairy and beef cattle, before being slaughtered/culled. It is important to note however, that the period during which the animals are subjected to these risks varies between production goals, production stages and the management goals of the farm (Figure 1A). Veal calves (from both dairy and beef herd origin) are usually slaughtered at the age of 6-11 months, depending on whether they are used to produce white or rose veal (but see culling of ‘surplus’/bobby dairy calves within the first days of life [43]). Calves raised for red-meat (from both dairy and beef herds) are usually slaughtered at 12-36 months, while replacement cows, which are used in both types of systems to produce both milk and calves, will commonly be slaughtered at an earlier age in dairy herds than beef herds. In the US, for example, dairy cows are slaughtered at about 5 years of age (after 2.5-3 lactations), while beef cows are slaughtered at 7-12 years of age [44]. The production of milk, like beef production, involves killing of animals, yet in many cases, this happens at a younger age in the dairy industry. Factoring the intensity of welfare compromise in the life time expectancy (i.e. duration) of these animals, could deepen our understanding of the overall welfare impact. However, it is beyond the scope of this paper. Here and throughout the manuscript, we avoid making any ethical claims about sanctity of life. Also, we do not aim to value the amount of suffering per animal in relation to the amount of animal protein (or calorie [45]). Our aim was more straightforward, to assess, based on expert opinion, the welfare of animals born to common dairy and beef herds, until slaughtered/culled.

At a time of rising public concern for the welfare of animals [46] and awareness of the impact of our diets on the environment [47], it is important that research priorities and dietary choices are aligned with the areas where welfare problems are most apparent. We call for further comparisons of bovine welfare in less common dairy and beef production systems that were not covered here (e.g. organic systems, dam rearing systems, where dairy calves are kept together with their dam), preferably using animal-based welfare assessment on farm. It would also be valuable to obtain representative overview of the prevalence of these different systems across countries. In addition, we encourage a similar expert comparison of other farm species that, like dairy cows, are used for “more than their meat”; e.g. laying hens could be compared with broilers to see if the findings from this study apply in other areas of animal agriculture.

## Conclusion

Cattle welfare experts rated dairy cattle as significantly/substantially more likely to experience negative welfare than beef cattle in the most common housing systems selected by the experts. The underpinning reasons for these evaluations were not explored with experts but are proposed to enable testable predictions for future research. Exploring the alignment between consumer choices and independent evaluations of animal welfare may encourage a more sustainable and responsible food production and consumption.

## Author contributions

RM, LG, MBMB and CJN designed research; RM performed research; LG and RM analyzed data; RM, LG, MBMB, CJN and JAW wrote the paper.

## Supporting information

### Supplementary text

#### PCA, 12 animal welfare statements

The PCA on the residuals of the normalised values of the 12 welfare statements resulted in a first principal component (PC) that explained 50.3% of the overall variability, which was clearly more than any other PC (S2 Table, S4 Fig.).

As all the original variables loaded positively on this first PC, it can be interpreted accordingly as the overall welfare risk summarising all the 12 original welfare statements. The highest loads were reached by the risk of restricted ability to express other normal behaviours, of restricted movement, of restricted ability to express social behaviours, of discomfort when resting, of experiencing negative affective states and of restricted experience of positive emotions (S2 Table). The beef versus dairy origin was well separated along this first PC (Fig. 1 in main text; S4 Fig.).

All additional PCs explained a much smaller proportion of the variance and were mostly difficult to interpret. Due to their limited contribution to the overall variability in the data, no attempt was made at their interpretation and they were not further considered for analyses.

#### PCAs, information on the housing systems

The analyses of the information on the housing systems showed that beef versus dairy origin can be well separated along the first PC in all the animal categories (S3 Table, S5 Fig., a-d).

For the calves raised for red-meat, the housing systems could be described by two PCs, of which the first was composed of access to pasture, access to dam and suckling from dam during the first months of life and not being transported during the first 6 months of life (“extensitivity” which may be closely related to available space), whereas the second concerned being slaughtered before or after 18 months of age (S3 Table). Beef and dairy systems seemed to differ mostly by the beef systems being more extensive (S5a Fig.). For the calves raised for replacement, the results were very similar with the exception that the transportation loaded on the second PC (S3 Table, S5b Fig.) and that, due to their purpose, they were not slaughtered by default.

The housing of beef and dairy cows seemed to vary mostly in the extent that access to the outdoors was provided (S3 Table, S5c Fig.). Here, beef cows seemed to have access to the outdoors more commonly than dairy cows and dairy cows ran a higher risk of being culled early.

The housing of veal calves seemed to vary the most in respect to the type of flooring provided with bedding material and access to pasture versus housing on slatted floors loading heavily on the first PC (S3 Table, S5d Fig.). The second PC was a contrast of the provision of (additional) solid feed and being slaughtered at a young age.

**S1 Figure.**
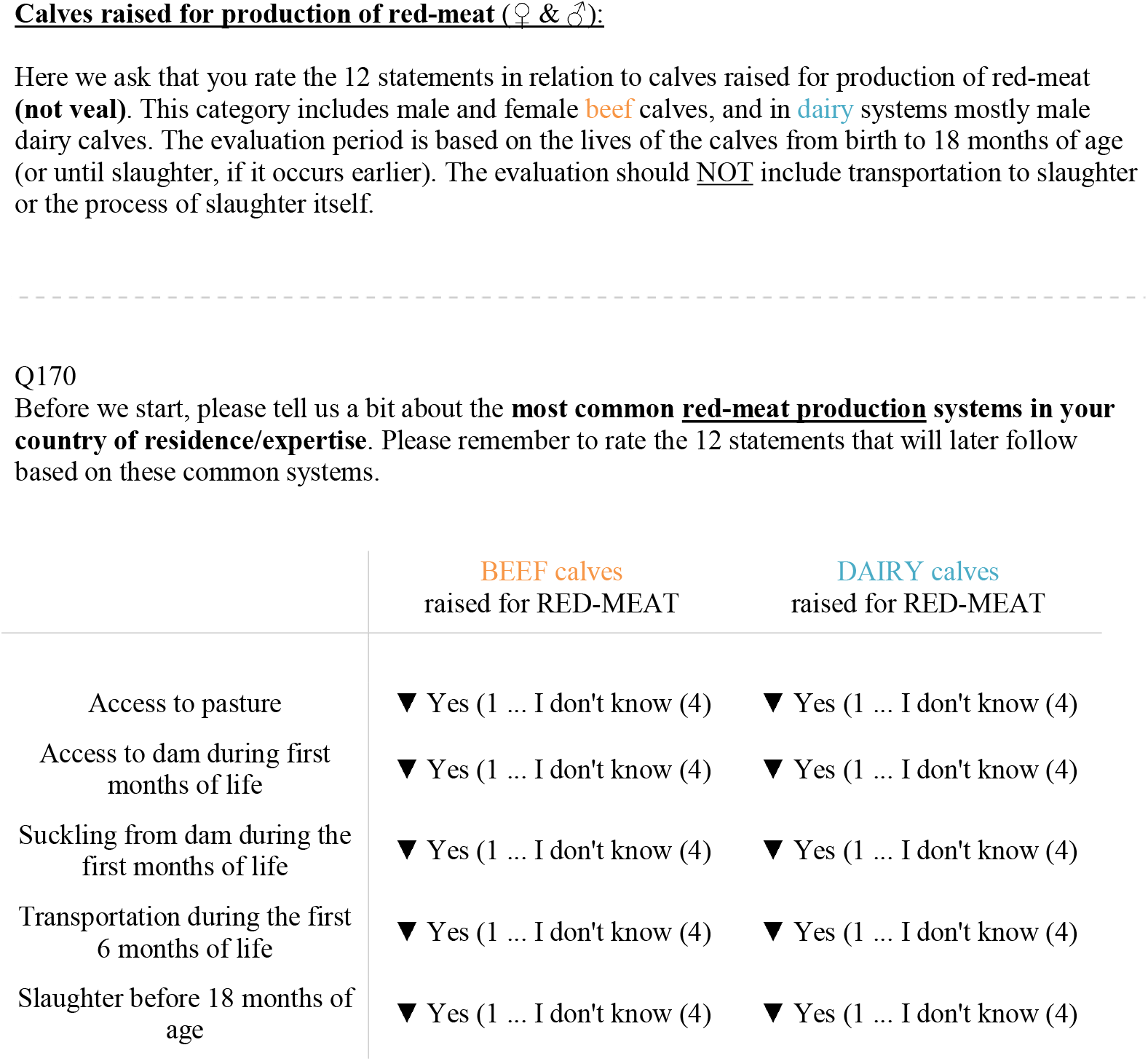
Illustration of part 3 of the survey. Characterization of the common production systems. Similar pages were shown for the three other animal categories.

**S2 Figure.**
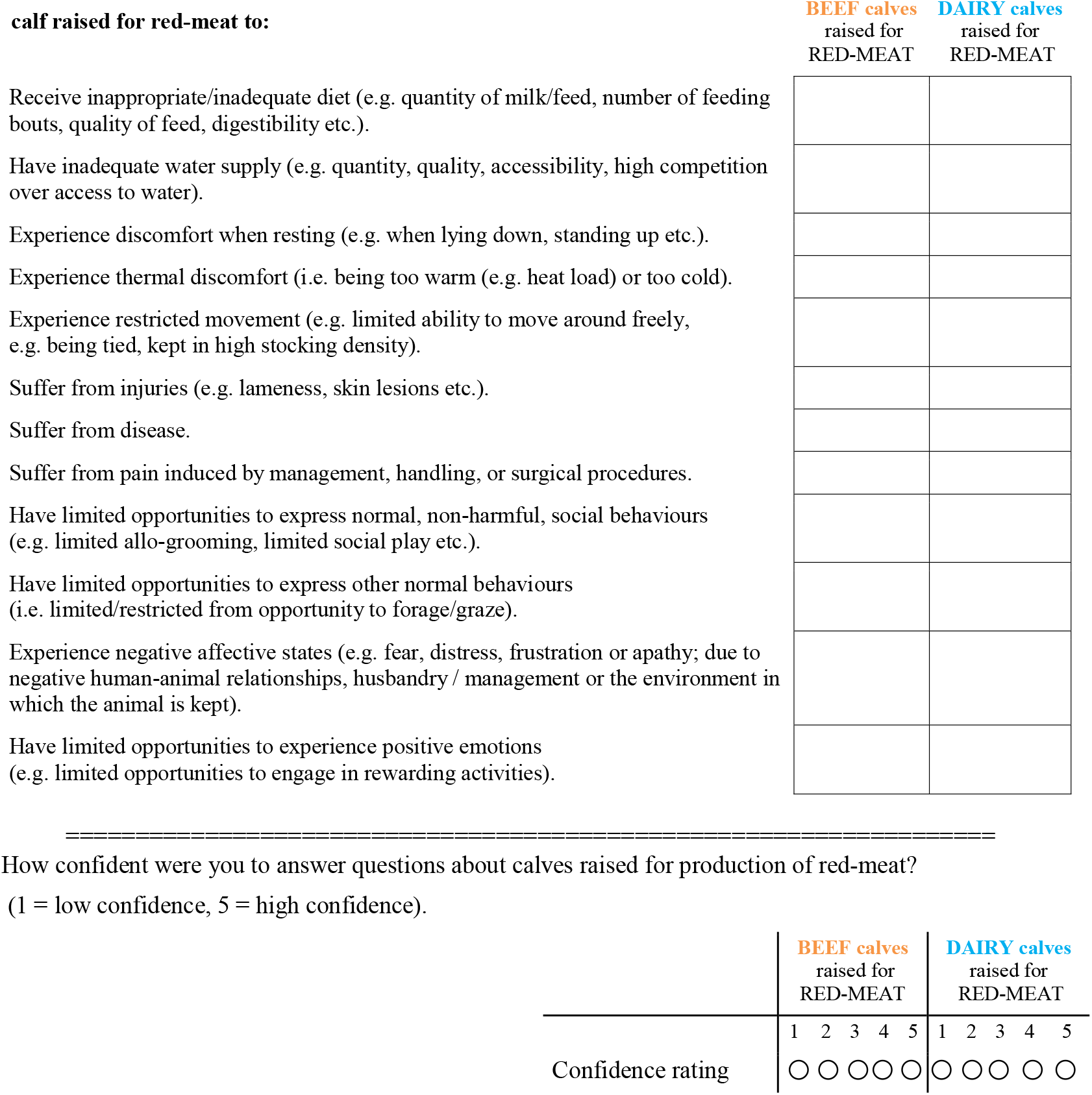
Illustration of part 3 of the survey. Please rate the likelihood of the statements below on a scale of 1 (very low) to 7 (very high). Notice that the statements are built in such a way that the higher the likelihood ratings are, the higher is the welfare-risk for the animal.

**S3 Figure.**
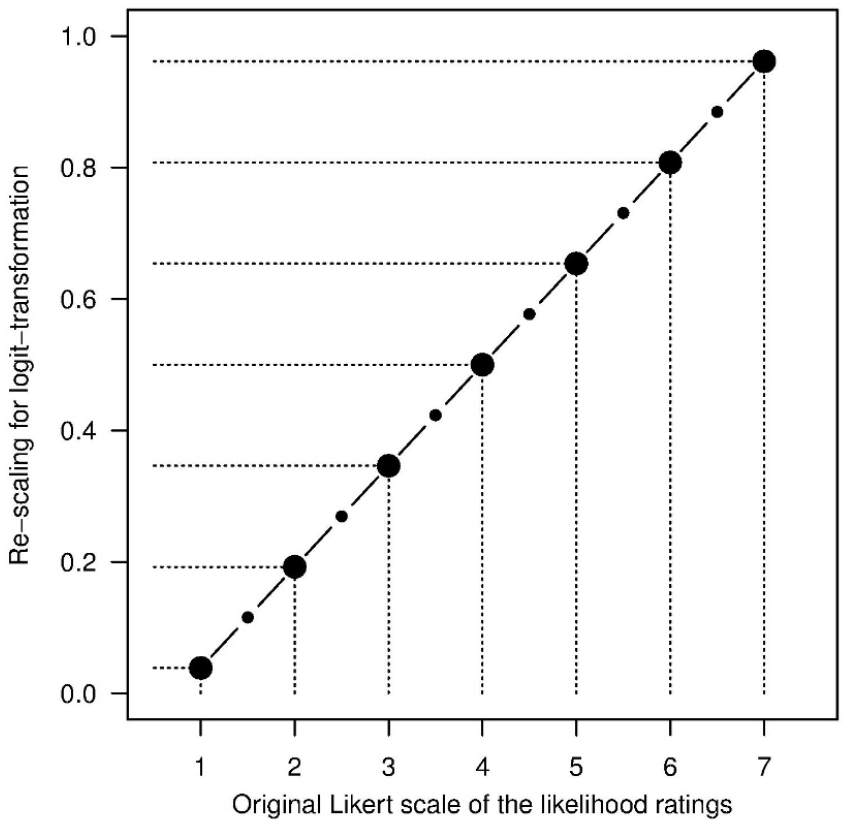
Re-scaling the likelihood rating. Re-scaling the original likelihood rating on a Likert scale (X-axis) to a proportion scale (Y-axis) that was, then, logit-transformed

**S4 Figure.**
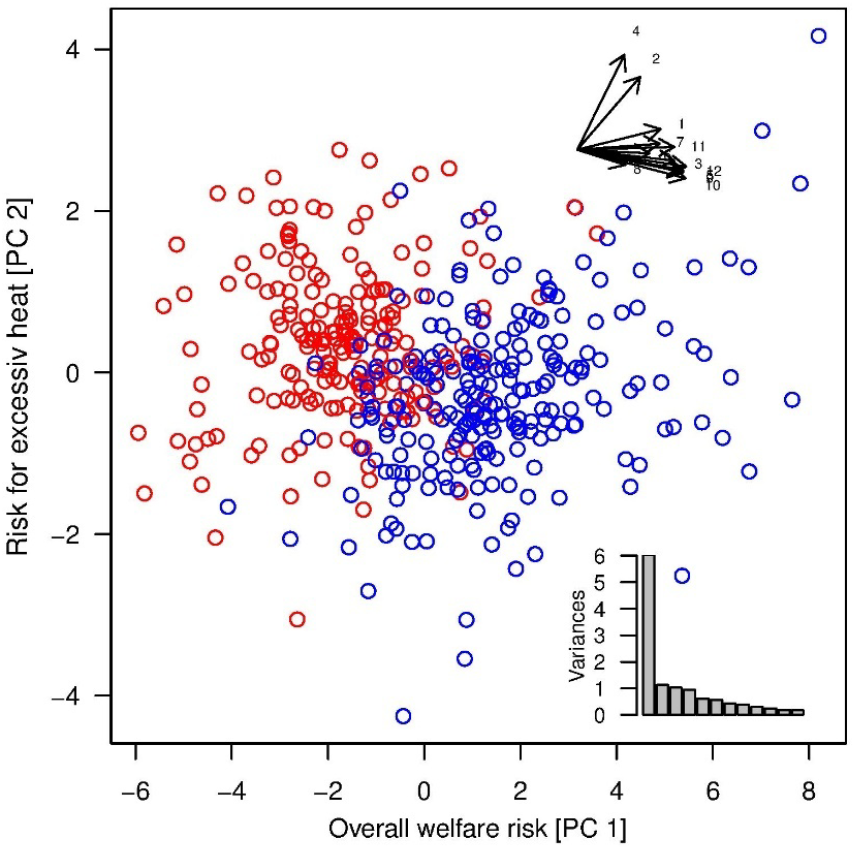
Result of the PCA on residuals of the normalised values. All observations are shown in the space spanned by the 1^st^ and 2^nd^ PC (beef origin in red and dairy origin in blue). The directions of the twelve welfare statements is given in the top-inset. The relative contribution of the 12 PCs to the overall variance is given in the bottom-inset. See Table 1 for the raw scores of the original questions and S2 Table for the loadings of the original variables on the PCs.

**S5 Figure.**
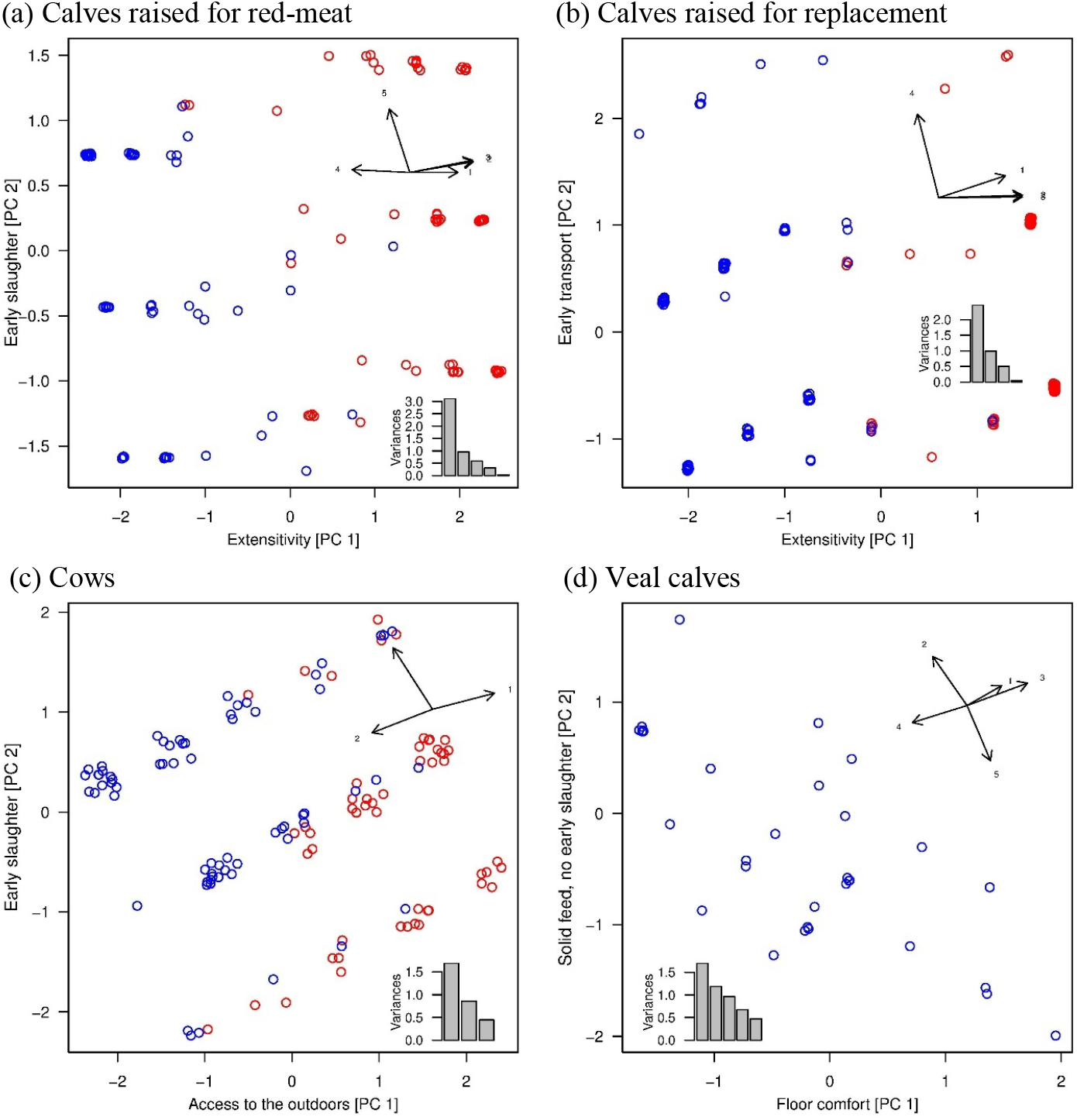
Result of the PCAs for the information on the housing systems. In each diagram, all observations are shown in the space spanned by the 1^st^ and 2^nd^ PC (beef origin in red and dairy origin in blue). Points are jittered to show the number of points with the same values on the PCs. The directions of the original questions (S3 Table) is given in the top-inset. The relative contribution of the PCs to the overall variance is given in the bottom-inset.

**S1 Table.**
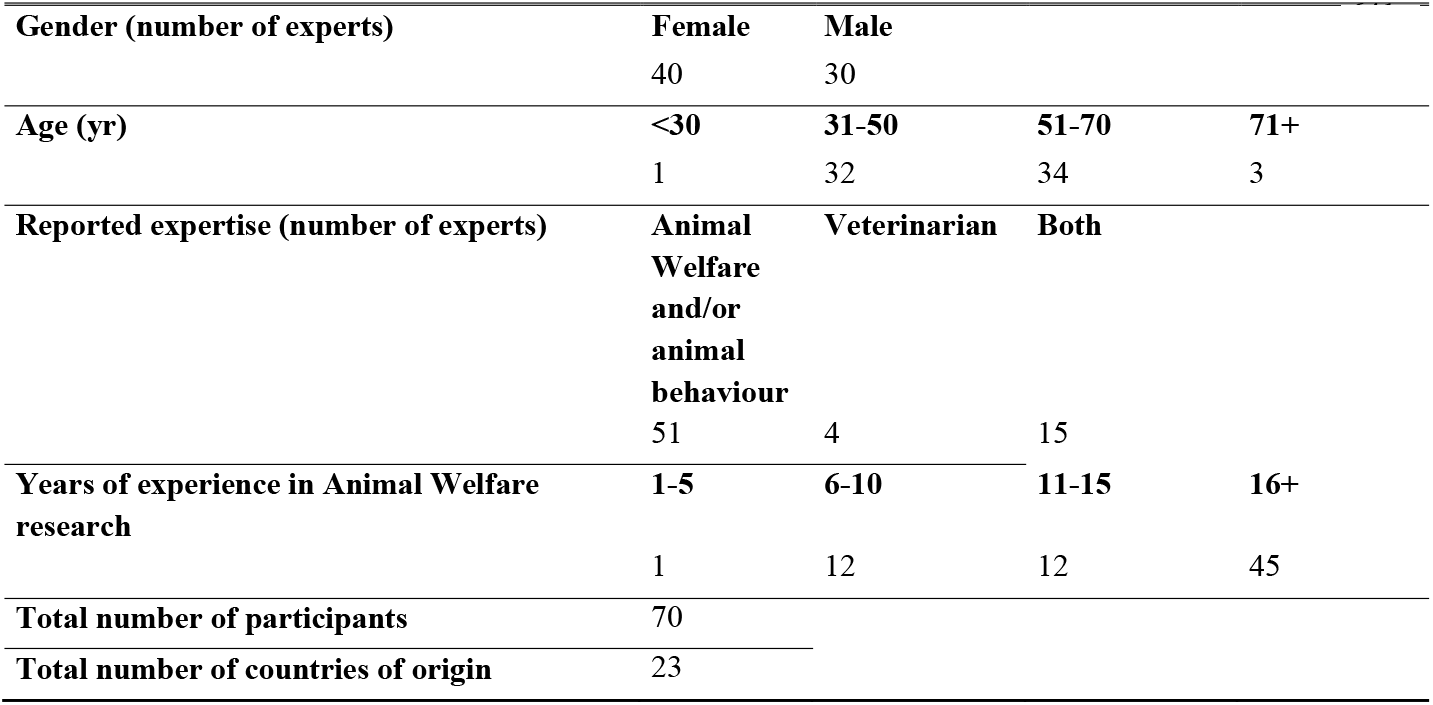
Expert characteristics.

**S2 Table.**
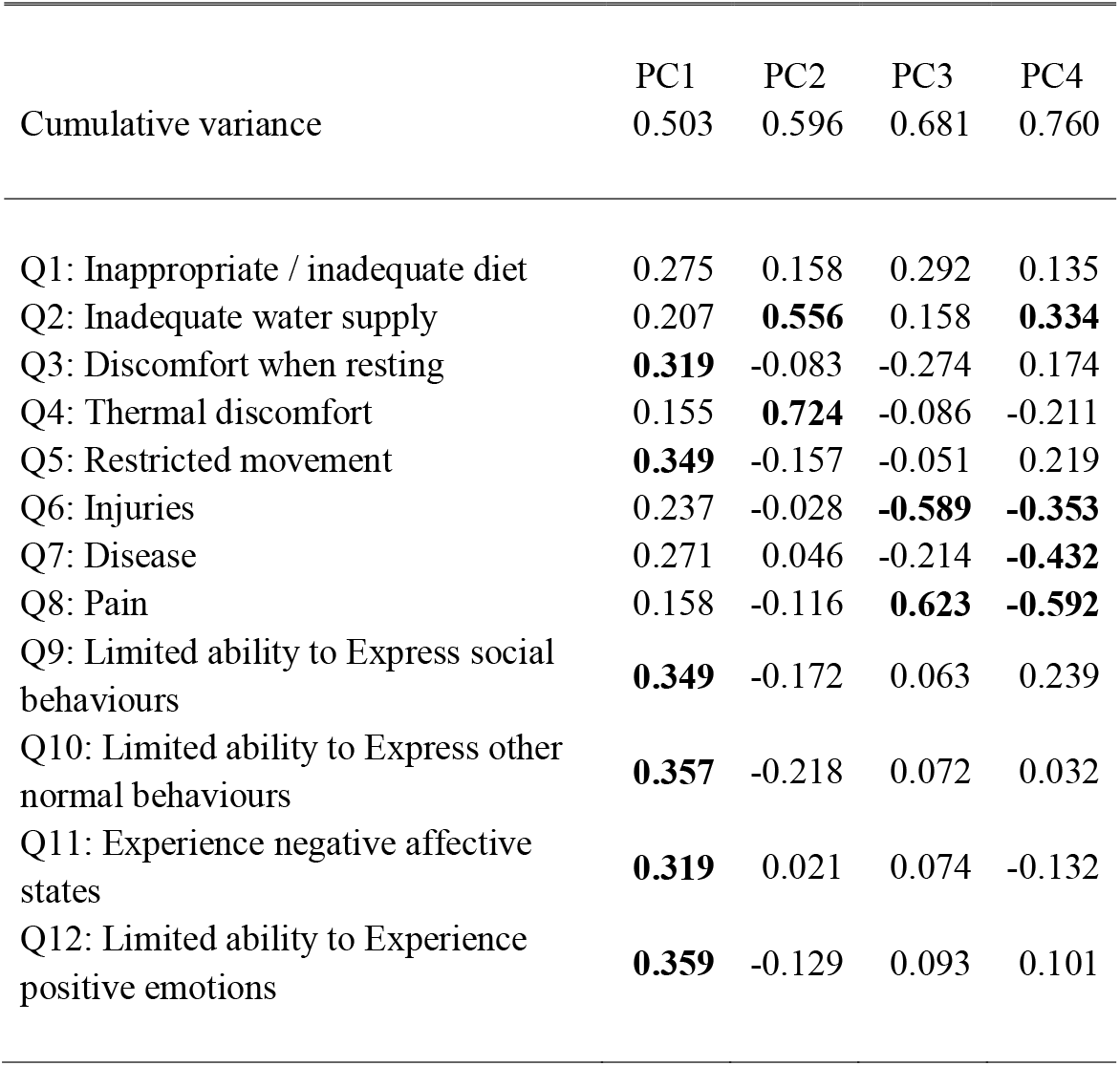
Loadings of the normalised values of the 12 welfare statements on the principle components and cumulative variance explained by the principle components. (Components larger than 0.3 are marked in bold).

|  | PC1 | PC2 | PC3 | PC4 |
| --- | --- | --- | --- | --- |
| Cumulative variance | 0.503 | 0.596 | 0.681 | 0.760 |
| Q1: Inappropriate / inadequate diet | 0.275 | 0.158 | 0.292 | 0.135 |
| Q2: Inadequate water supply | 0.207 | **0.556** | 0.158 | **0.334** |
| Q3: Discomfort when resting | **0.319** | -0.083 | -0.274 | 0.174 |
| Q4: Thermal discomfort | 0.155 | **0.724** | -0.086 | -0.211 |
| Q5: Restricted movement | **0.349** | -0.157 | -0.051 | 0.219 |
| Q6: Injuries | 0.237 | -0.028 | **-0.589** | **-0.353** |
| Q7: Disease | 0.271 | 0.046 | -0.214 | **-0.432** |
| Q8: Pain | 0.158 | -0.116 | **0.623** | **-0.592** |
| Q9: Limited ability to Express social behaviours | **0.349** | -0.172 | 0.063 | 0.239 |
| Q10: Limited ability to Express other normal behaviours | **0.357** | -0.218 | 0.072 | 0.032 |
| Q11: Experience negative affective states | **0.319** | 0.021 | 0.074 | -0.132 |
| Q12: Limited ability to Experience positive emotions | **0.359** | -0.129 | 0.093 | 0.101 |

**S3 Table.** Loadings of the questions on housing conditions of calves kept for the production of red meat, of calves kept as replacement animals, of cows kept for dairy or meat production, and of veal calves on the principle components and cumulative variance explained by the first two principle components each. (Components larger than 0.3 are marked in bold).

|  | Calves kept for the<br>production of red<br>meat |  |  | Calves kept as<br>replacement animals |  |  | Cows kept for dairy<br>or meat production |  |  | Veal calves |  |  |
| --- | --- | --- | --- | --- | --- | --- | --- | --- | --- | --- | --- | --- |
|  |  | PC1 | PC2 |  | PC1 | PC2 |  | PC1 | PC2 |  | PC1 | PC2 |
| Cumulative variance |  | 0.624 | 0.816 |  | 0.619 | 0.865 |  | 0.566 | 0.852 |  | 0.341 | 0.579 |
| Access to pasture | Q1 | **0.407** | 0.002 | Q1 | **0.486** | 0.256 | Q1 | **0.652** | 0.237 | Q1 | **0.354** | 0.244 |
| Access to dam during first months of life | Q2 | **0.536** | 0.164 | Q2 | **0.608** | 0.030 |  |  |  |  |  |  |
| Suckling from dam during the first months of life | Q3 | **0.530** | 0.187 | Q3 | **0.608** | 0.010 |  |  |  |  |  |  |
| Provided with solid feed, in addition to milk/milk<br>replacer (from age of 2 w) |  |  |  |  |  |  |  |  |  | Q2 | **-0.350** | **0.601** |
| Provided with bedding material (from age of 2 w) |  |  |  |  |  |  |  |  |  | Q3 | **0.618** | 0.275 |
| Slatted floor from the age of 8 w onwards |  |  |  |  |  |  |  |  |  | Q4 | **-0.559** | -0.209 |
| Indoor housing |  |  |  |  |  |  | Q2 | **-0.635** | -0.348 |  |  |  |
| Transportation during first 6 mo of life | Q4 | **-0.487** | 0.043 | Q4 | -0.154 | **0.966** |  |  |  |  |  |  |
| Slaughtered before 8 mo of age |  |  |  |  |  |  |  |  |  | Q5 | 0.240 | **-0.678** |
| Slaughter before 18 mo of age | Q5 | -0.173 | **0.968** |  |  |  |  |  |  |  |  |  |
| Slaughter age below 6 y |  |  |  |  |  |  | Q3 | **-0.414** | **0.907** |  |  |  |

